# “Alkyladenine DNA glycosylase associates with transcription elongation to coordinate DNA repair with gene expression”

**DOI:** 10.1101/802983

**Authors:** Nicola P. Montaldo, Diana L. Bordin, Alessandro Brambilla, Marcel Rösinger, Sarah L. Fordyce Martin, Karine Øian Bjørås, Stefano Bradamante, Per Arne Aas, Antonia Furrer, Lene C. Olsen, Marit Otterlei, Pål Sætrom, Magnar Bjørås, Leona D. Samson, Barbara van Loon

## Abstract

Base excision repair (BER) initiated by alkyladenine DNA glycosylase (AAG; aka MPG) is essential for removal of aberrantly methylated DNA bases. Genome instability and accumulation of aberrant bases accompany multiple diseases including cancer and neurological disorders. While BER is well studied on naked DNA, it remains unclear how BER efficiently operates on chromatin. Here we show that AAG binds to chromatin and forms complex with active RNA polymerase (pol) II. This occurs through direct interaction with Elongator and results in transcriptional co-regulation. Importantly, at co-regulated genes aberrantly methylated bases accumulate towards 3’end, in regions enriched for BER enzymes AAG and APE1, Elongator and active RNA pol II. Active transcription and functional Elongator are further crucial to ensure efficient BER, by promoting AAG and APE1 chromatin recruitment. Our findings provide novel insights to maintaining genome stability in actively transcribing chromatin, and reveal roles of aberrantly methylated bases in regulation of gene expression.

## Introduction

Thousands of DNA base lesions are generated in each cell of our body every day as a consequence of exposure to endogenous and exogenous DNA damaging agents^1^. Aberrantly methylated bases account for a large proportion of those lesions. Estimated steady-state levels of the most frequent aberrantly methylated bases: 7-methylguanine (7meG) and 3-methyladenine (3meA), are 6,000 and 1,200 lesions per mammalian cell per day respectively^2^. Several lines of evidence strongly indicate that accumulation of aberrant DNA bases and genome instability accompany major human diseases like cancer, inflammatory diseases and neurological conditions^3,4^.

Base excision repair (BER) is a highly efficient mechanism for the removal of aberrant DNA bases. This multi-step repair process is initiated by substrate-specific DNA glycosylases that recognize and remove aberrant bases^3,5^. The resulting abasic (AP) site is processed by the apurinic/apyrimidinic endonuclease 1 (APE1) that generates a single-strand break (SSB), which is subsequently filled by DNA polymerase (pol) β, and the resulting nick sealed by DNA ligase III/XRCC1^6^. The BER specificity is determined by the type of DNA glycosylase that initiates the pathway^7^. Alkyladenine DNA glycosylase (AAG, aka. N-methylpurine-DNA glycosylase, MPG) is the major mammalian DNA glycosylase responsible for the removal of aberrantly methylated bases 7meG and 3meA^6,7^. It participates in both nuclear and mitochondrial DNA repair^8^. Much of our current knowledge about the different BER steps derives from *in vitro* studies using naked DNA as a substrate. However, eukaryotic cells need to repair DNA in the context of chromatin. Recent work demonstrated that the activity of human AAG, and of several other DNA glycosylases, is strongly impaired in the context of nucleosomes^9^. AAG activity is dramatically inhibited (84% to 100%) when the aberrant base is positioned mid-way or directly faces the surface of the histone octamer. Furthermore, the presence of nucleosomes was shown to significantly impair the activity of the downstream BER proteins APE1, DNA pol β and DNA ligase III/XRCC1^10-12^. It was proposed that BER could overcome the inhibitory effect of nucleosomes by associating with processes that involve chromatin reorganization, such as replication and transcription^13-15^. This idea is further supported by the notion that BER preferentially occurs on the transcribed strand^16,17^. However, firm evidence demonstrating that BER associates with the transcription is still lacking. AAG could potentially be associated with the transcription through interaction with different transcription factors. It was shown that AAG forms a complex with a transcriptional repressor methylated DNA-binding domain 1 (MBD1) and that this complex could inhibit the transcription^18^. Furthermore, AAG was demonstrated to bind to the estrogen receptor (ER) α, which in turn stimulates AAG glycosylase activity^19^ and could potentially result in repression of the ERα-responsive genes. Interestingly, recent work in mice focusing on DNA glycosylases responsible for the repair of oxidized bases, similarly, suggested the role of glycosylases in regulation of ERα-gene expression^20^. Taken together, these findings indicate that coupling of AAG-initiated BER to transcription could enable more efficient repair and that AAG potentially influences gene expression.

Recent extensive protein interaction study^21^ predicted that AAG forms a complex with the active transcription machinery through a possible interaction with Elongator complex. Elongator is a highly conserved complex that participates in several pathways, including facilitation of transcription elongation by interacting with hyperphosphorylated RNA pol II^22,23^. It is composed of six subunits ELP1 to ELP6 organized in a dodecamer^24,25^. Every subunit is required for the complex to function. ELP1 (aka. IKBKAP) is the largest subunit of the complex and provides a scaffolding function^26^. ELP3 has histone acetyltransferases (HAT) activity and acetylates histone H3^27,28^. Mutations in human *ELP1*, and consequently impaired Elongator function cause neurodevelopmental disorder, and lead to reduced expression of Elongator-dependent genes in patient-derived cells^29^. Interestingly, loss of Elp1 in yeast causes hypersensitivity to methyl methanesulfonate (MMS) and hydroxyurea^30^. While the predicted association between AAG and Elongator complex suggests that AAG-initiated BER may be coordinated with transcription elongation, the interaction remains yet to be confirmed and its relevance explored. Taken together, several studies suggest that AAG could form a complex with different transcriptional components, however whether and where these interactions take place in the context of chromatin, as well as the extent to which they facilitate AAG-initiated BER and regulate gene expression remains unknown.

In this study we show that the majority of cellular AAG localizes at chromatin and is engaged in a complex with actively transcribing RNA pol II. This occurs through direct interaction with the ELP1 subunit of transcriptional Elongator complex, and is to our knowledge the first evidence of BER association with active transcription. RNA-sequencing experiments further show that AAG and ELP1 co-regulate genes, which are primarily repressed by AAG and stimulated by ELP1. Notably, at co-regulated genes endogenous aberrantly methylated DNA bases accumulate towards 3’end, in regions co-occupied by ELP1, elongating RNA pol II, and BER enzymes AAG and APE1. Chromatin recruitment of the BER enzymes is strongly dependent on Elongator presence and active transcription, since ELP1 loss as well as transcription inhibition cause globally reduced AAG and APE1 binding. Importantly, functional Elongator and active transcription are needed to ensure efficient AAG-initiated BER, and their inactivation results in impaired repair and significant accumulation of AAG substrates genome wide. Based on our findings, we propose that AAG, in concert with Elongator complex and active transcription, coordinates repair of aberrantly methylated DNA bases with regulation of gene expression.

## Results

### AAG DNA glycosylase associates with active transcription to regulate gene expression

AAG predominantly localizes to the nucleus where it recognizes and removes aberrantly methylated DNA bases. However, it remains unknown how AAG initiates repair in the context of chromatin, and in which specific regions of the genome the repair takes place. To determine which portion of nuclear AAG is bound to the chromatin and could actively participate in the repair of genomic DNA, HEK293T cells were fractionated. The amount of AAG detected in the chromatin fraction (CF) was comparable to the total AAG fraction (TF), while only low levels of AAG were detected in the soluble fraction (SF) (Fig. 1a, b), thus indicating that the majority of cellular AAG is chromatin bound. In order to efficiently repair aberrant bases in chromatin context, DNA glycosylases were suggested to associate with processes that involve chromatin reorganization, such as transcription^13^. To determine if AAG engages with the active transcription, immunoprecipitation (IP) was performed and AAG-specific cellular complexes analyzed. Interestingly, the IP analysis indicated that AAG forms a complex with hyperphosphorylated RNA pol II (Fig. 1c). This is to our knowledge the first evidence of a DNA glycosylase forming a complex with the core transcriptional RNA pol II. To determine regions in which transcription is influenced by AAG presence, we generated HEK293T cells lacking AAG (AAG^−/-^) using the CRISPR-Cas9 engineering approach (Fig. 1d), and subjected them to RNA sequencing (Fig. 1e). Comparison of wild type (WT) and AAG^−/-^ transcriptomes revealed that, at ≥1.5-fold change and an FDR≤0.1, 1,045 genes were differentially expressed in AAG^−/-^ cells. The subsequent DAVID gene ontology (GO) enrichment analysis^31^ revealed that the majority of differentially expressed genes (DEGs) in AAG^−/-^ cells belonged to neurogenesis and nervous system development processes (Fig. 1f). These findings are further supported by analysis of DEGs in HAP1 AAG^−/-^ cells, which similarly determined the neurodevelopmental processes as the most significantly enriched (Supplementary figure 1). Subsequent separation of DEGs based on the expression change, revealed that the large portion of genes affected by the loss of AAG are upregulated (Fig. 1g). These upregulated genes were the main drivers of the GO segregating in the nervous system development and neurogenesis processes (Fig. 1h), while the down-regulated genes produced no significant GO terms (Fig. 1i). Taken together these findings suggest that AAG binds to chromatin, forms a complex with the active transcription and regulates gene expression.

**Figure 1.**
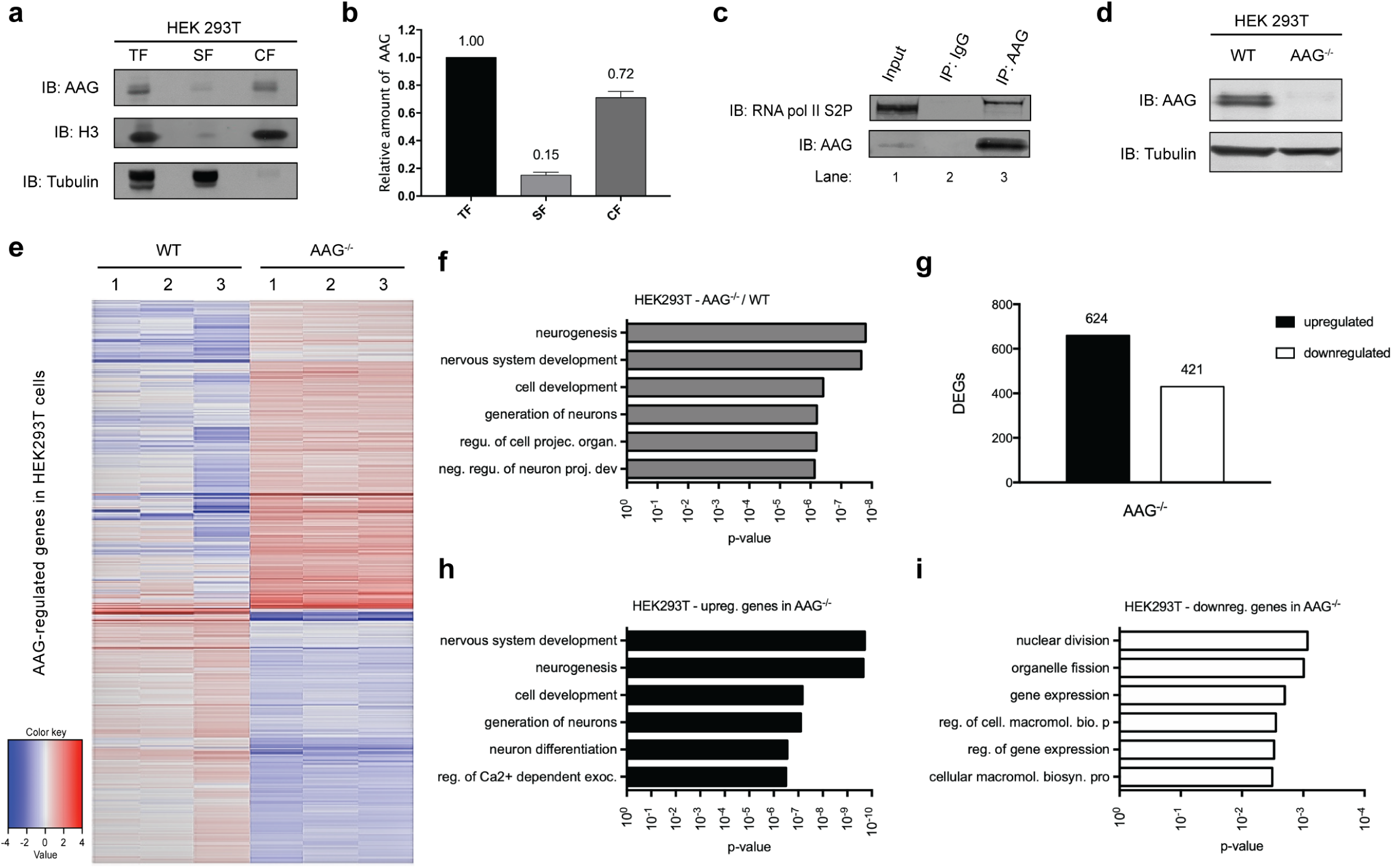
AAG DNA glycosylase associates with active transcription and regulates gene expression. **a** Immunoblot analysis of chromatin fractionation assay indicating AAG distribution in total fraction (TF), soluble supernatant fraction (SF) and chromatin fraction (CF). Histone H3 and α-tubulin served as controls. **b** Quantification of three biologically independent experiments as the one depicted in (a); error bars represent SEM. **c** Immunoprecipitation of AAG from HEK293T whole cell extracts showing the interaction with RNA polymerase II phosphorylated at Serine 2 (S2P) of CTD repeat. **d** Immunoblot analysis of HEK293T WT and AAG^−/-^ whole cell extracts generated by CRISPR-Cas9 technology. **e** Heat map of the expression levels of AAG-regulated genes in HEK293T cells. **f** Top six biological processes (BP) gene ontology (GO) terms as determined by the Database for Annotation, Visualization and Integrated Discovery (DAVID) for genes dysregulated in AAG^−/-^ HEK293T cells when compared to WT. **g** Up- and down-regulated differentially expressed genes (DEGs) in AAG^−/-^ HEK293T cells. **h** and **i** Top six BP GO terms as determined by DAVID for genes upregulated (h) and downregulated (i) genes.

### AAG directly interacts with the transcriptional Elongator complex

AAG-initiated BER can be linked to transcription through direct association with RNA pol II or by binding other components of the transcription machinery. To identify proteins that form a complex with AAG, liquid chromatography–tandem mass spectrometry (LC/MS-MS) analysis was performed in HEK293T cells expressing FLAG-tagged AAG, either untreated or exposed to the alkylating agent MMS (Fig. 2a). The LC/MS-MS analysis revealed that the ELP1, 2 and 3, all components of the transcriptional Elongator complex, were among the most enriched proteins (Fig. 2a, b). MMS exposure did not appear to affect AAG-Elongator complex formation, since a comparable number of peptide spectra were detected in the untreated and treated samples (Fig. 2b, Supplementary table 1). Subsequent IP experiments showed that, similar to the exogenous FLAG-AAG, the endogenous AAG forms a complex with ELP1 independently of the MMS treatment (Fig. 2c). Further, the separation of protein complexes present in the HeLa whole cell extract (WCE) revealed that the main Elongator subunits co-elute with AAG (Supplementary figure 2a). Since both AAG and Elongator are DNA binding proteins, we next examined whether the complex formation is dependent on the presence of nucleic acids. Addition of different nucleases (DNaseI, MNase and RNaseI) during either AAG or ELP1 IP did not influence the levels of co-precipitated proteins (Supplementary figure 2b, c). These findings thus suggest that in human cells endogenous AAG forms a complex with Elongator, independently of the presence of nucleic acids and exposure to genotoxic stress. To examine if AAG binds directly to Elongator, we purified recombinant full-length (fl) AAG, AAG lacking 80 N-terminal amino acids (Δ80) and the FLAG-tagged ELP1 (Supplementary figure 2d-g). ELP1 was targeted, since it was the predominant Elongator subunit detected in the complex with AAG (Fig. 2a, b). Incubation of recombinant proteins and subsequent IP analysis indicated that both purified fl and Δ80 AAG interact with FLAG–ELP1, although the amount of precipitated Elongator was markedly reduced with Δ80 AAG (Fig. 2d). These findings indicate that AAG directly interacts with the ELP1 subunit of the transcriptional Elongator complex, predominantly through unstructured N-terminal domain.

**Figure 2.**
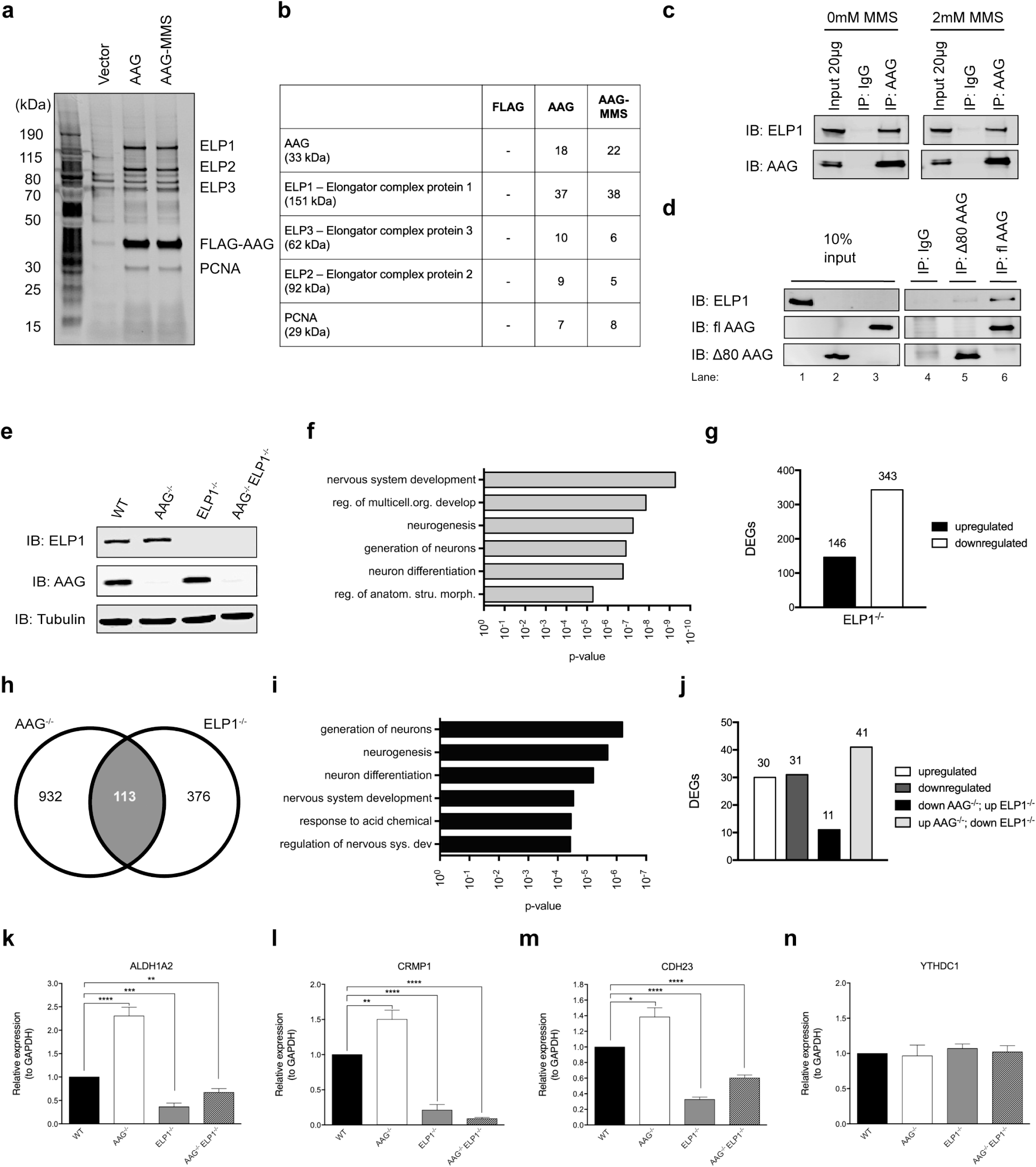
AAG directly interacts with ELP1 subunit of transcriptional Elongator complex and the two co-regulate gene expression. **a** Silver staining analysis of FLAG-AAG IP of samples subjected to LC/MS-MS analysis. **b** AAG interacting partners identified via LC/MS-MS analysis. The number indicates the exclusive unique spectra counts assigned to a protein. See also Supplementary table 1. **c** Immunoprecipitation (IP) of AAG from HEK293T whole cell extracts (WCEs) treated or untreated with MMS. **d** Co-IP of purified FLAG-ELP1 with AAG full length (fl AAG) or AAG lacking 80 N-terminal amino acids (Δ80 AAG). **e** Immunoblot of HEK293T whole cell extracts (WCEs) from WT, AAG^−/-^, ELP1^−/-^ and AAG^−/-^ELP1^−/-^ cell lines generated via CRISPR-Cas9 technology. **f** Top six biological processes (BP) gene ontology (GO) terms as determined by DAVID for genes dysregulated in HEK293T ELP1^−/-^ cells when compared to WT cells. **g** Downregulated and upregulated differentially expressed genes (DEGs) in HEK293T ELP1^−/-^ cells. **h** Venn diagrams of AAG- and ELP1-regulated genes in HEK293T cells. **i** Top six BP GO terms as determined by DAVID of genes co-regulated by AAG and ELP1 in HEK293T cells. **j** Directionality of DEGs regulated by AAG and ELP1 in HEK293T cells. **k-n** Relative mRNA levels of *ALDH1A2* (k), *CRMP1* (l), *CDH23* (m) and *YTHDC1* (negative control) (n) genes in HEK293T WT, AAG^−/-^, ELP1^−/-^ and AAG^−/-^ELP1^−/-^ cells. Error bars, represent SEM from at least three independent experiments. *p<0.05, **p<0.005, ***p<0.0005, two-tailed Student’s t-test; NS, not significant.

### AAG and Elongator co-regulate gene expression

To identify genes and regions at which AAG and Elongator have the most significant impact we generated, in addition to AAG^−/-^ cells, the HEK293T cells lacking ELP1 (ELP1^−/-^), and both AAG and ELP1 (AAG^−/-^ELP1^−/-^) (Fig. 2e), via the CRISPR-Cas9 approach. Subsequent RNA sequencing analysis of ELP1^−/-^ cells showed that loss of ELP1 results in ≥2-fold altered expression of 489 genes, with FDR≤0.1. Similar to AAG^−/-^ cells (Fig. 1f), DEGs in ELP1^−/-^ cells belonged to the nervous system development processes (Fig. 2f). The majority of the DEGs were downregulated in ELP1^−/-^ cells (343 out of 489) (Fig. 2g), thus supporting previous observation that Elongator promotes transcription^29^. Comparison of genes differentially expressed in AAG^−/-^ and ELP1^−/-^ cells resulted in identification of 113 co-regulated genes (Fig. 2h) that predominantly clustered into processes involving generation of neurons and neurogenesis (Fig. 2i). Interestingly, the majority of co-regulated genes were upregulated in AAG^−/-^ and downregulated in ELP1^−/-^ cells (Fig. 2j), thus following the general direction of all DEGs in these two cell lines (Fig. 1g and 2g). The RNA sequencing results were confirmed by qPCR measurements of mRNA levels for multiple co-regulated genes belonging to the top identified biological processes presented in Fig. 2i. The mRNA levels of ALDH1A2, CRMP1, CDH23, SYT9, CDH4, NPTX2, NOVA2 and CDH22 were, as expected, significantly increased in AAG^−/-^, and decreased in ELP1^−/-^ cells (Fig. 2k-m and Supplementary figure 3), while the level of non-affected control YTHDC1 was unchanged in all tested cell lines (Fig. 2n). The observed increase in the expression of co-regulated genes was similar in two independent AAG^−/-^ clones (Supplementary figure 4). Notably, in AAG^−/-^ELP1^−/-^ cells, the co-regulated genes were significantly downregulated in comparison to WT levels, thus following the direction of ELP1^−/-^ cells (Fig. 2k-m and Supplementary figure 3). This suggests that at the transcriptional level Elongator acts upstream of AAG. In summary, these results provide evidence of genes co-regulated by the AAG DNA repair glycosylase and its interaction partner the Elongator complex.

**Figure 3.**
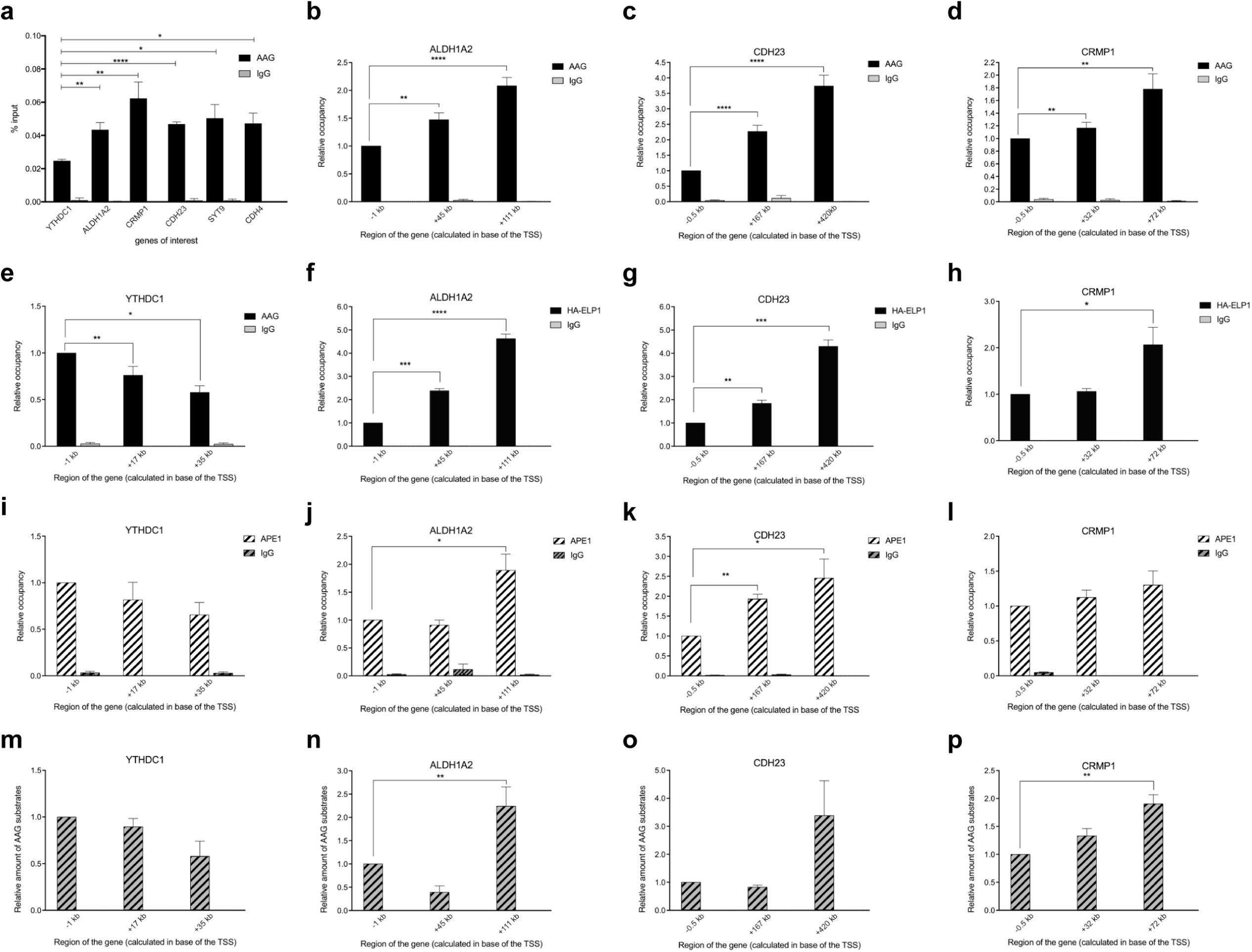
Elongator and components of AAG-initiated BER accumulate towards 3’end of co-regulated genes, in regions with high levels of aberrantly methylated AAG substrates. **a** AAG ChIP assays in HEK293T WT cells comparing % of input at gene bodies of unaffected gene (*YTHDC1*) and affected genes (*ALDH1A2, CRMP1, CDH23, SYT9, CDH4*). **b-e** ChIP assays showing relative AAG occupancy in AAG- and ELP1-dependent genes *ALDH1A2* (b), *CDH23* (c), *CRMP1* (d) and unaffected gene *YTHDC1* (e) in HEK293T WT cells. Error bars, SEM from at least three independent experiments. **f-h** ChIP assays showing relative occupancy of HA-ELP1 in AAG- and ELP1-dependent *ALDH1A2* (f), *CDH23* (g), *CRMP1* (h) genes in HEK293T WT cells with HA-ELP1. Error bars, SEM from two independent experiments. **i-l** ChIP assays showing relative APE1 occupancy in negative control *YTHDC1* gene (i), and AAG- and ELP1-dependent *ALDH1A2* (j), *CDH23* (k), *CRMP1* (l) genes in HEK293T WT cells. **m-p** qPCR DNA damage assay showing differences in distribution pattern of aberrant AAG substrates in unaffected gene Y*THDC1* (m) and genes regulated by AAG and ELP1: *ALDH1A2* (n), *CDH23* (o), *CRMP1* (p) in HEK293T WT cells. Error bars, SEM from at least three independent experiments. Values are shown as relative occupancy: % input of specific gene region relative to % input of promoter region. *p<0.05, **p<0.005, ***p<0.0005, two-tailed Student’s t-test.

**Figure 4.**
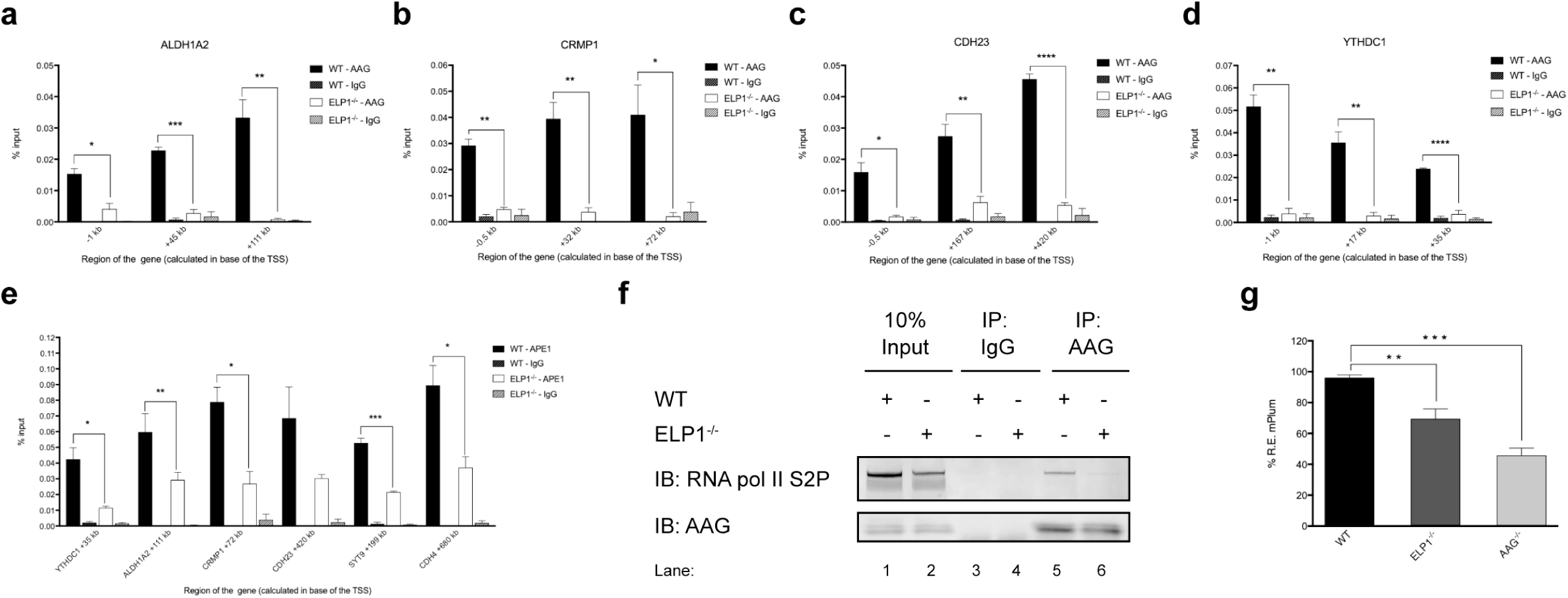
Loss of functional Elongator results in reduced AAG and APE1 chromatin recruitment, and impairs AAG-initiated repair. **a-d** ChIP-qPCR experiments comparing AAG occupancy at promoters and gene bodies of *ALDH1A2* (a), *CRMP1* (b), *CDH23* (c) and *YTHDC1* (d) genes in HEK293T WT and ELP1^−/-^ cells. **e** ChIP-qPCR experiments comparing APE1 occupancy at gene bodies of *YTHDC1, ALDH1A2, CRMP1, CDH23, SYT9* and *CDH4* genes in HEK293T WT and ELP1^−/-^ cells. Error bars, SEM from three independent experiments. *p<0.05, **p<0.005, ***p<0.0005, two-tailed Student’s t test. **f** Immunoprecipitation of AAG from HEK293T WT and Elp1^−/-^ whole cell extracts showing the interaction with RNA polymerase II phosphorylated at Serine 2 (S2P) of CTD repeat. **g** Measurement of AAG DNA glycosylase activity in HEK293T WT, AAG^−/-^ and ELP1^−/-^ on hypoxanthine (Hx)-containing plasmid by FM-HCR assays. Error bars represent the SEM calculated from at least four biological replicates; **p<0.005, ***p<0.0005, two-tailed Student’s t-test.

### AAG and Elongator accumulate toward 3’end of co-regulated genes in regions with high level of aberrantly methylated bases

To determine whether AAG and ELP1 localize at the co-regulated genes identified in Fig. 2, chromatin immunoprecipitation (ChIP) experiments were performed. AAG ChIP showed significant enrichment at the co-regulated genes (*ALDH1A2, CRMP1, CDH23, SYT9, CHD4*), when compared to *YTHDC1* unaffected control (Fig. 3a). Notably, the increase in AAG occupancy was most significant towards the 3’end of the co-regulated genes (*ALDH1A2, CRMP1, CDH23*) (Fig. 3b-d). This effect was specific for co-regulated genes and was not present at *YTHDC1* (Fig. 3e). To determine to which extent ELP1 distribution follows AAG occupancy, HEK293T cells with endogenously HA-tagged ELP1 were generated, using homologous recombination dependent CRISPR-Cas9 gene editing (Supplementary figure 5). Subsequent HA-ChIP experiments revealed that similar to the AAG distribution, and in line with its role in the transcription elongation, HA-ELP1 was significantly enriched towards the 3’end of the co-regulated genes (Fig. 3f-h). Importantly, the same distribution pattern was observed for RNA pol II phosphorylated at Serine 2 of the C-terminal domain (CTD) (RNA pol II S2P), which is the predominant form during transcription elongation (Supplementary figure 6). Further, to test if other BER enzymes co-occupy the same regions as AAG and ELP1, APE1 ChIP was performed. APE1 distribution closely resembled AAG and ELP1 localization at the co-regulated genes, with the highest occupancy detected towards the 3’end (Fig. 3i-l). Collectively, these results suggest that AAG-initiated BER associates with Elongator and transcription elongation, predominantly at the 3’end of the co-regulated genes. Since the main AAG function is to initiate BER, we next analyzed levels of aberrantly methylated AAG substrates along the co-regulated genes, using real-time qPCR based approach for quantification of aberrant DNA bases^32^. Interestingly, the distribution of endogenous aberrantly methylated AAG substrates closely followed AAG, APE1, ELP1 and RNA pol II occupancy, with the levels of aberrant bases being highest towards the 3’end of the co-regulated genes (Fig. 3m-p). Taken together these findings suggest that regions of co-regulated genes that are co-occupied by AAG-initiated BER and Elongator, have high levels of aberrant bases, thus indicating an interplay between the repair of DNA base lesions and transcription regulation.

**Figure 5.**
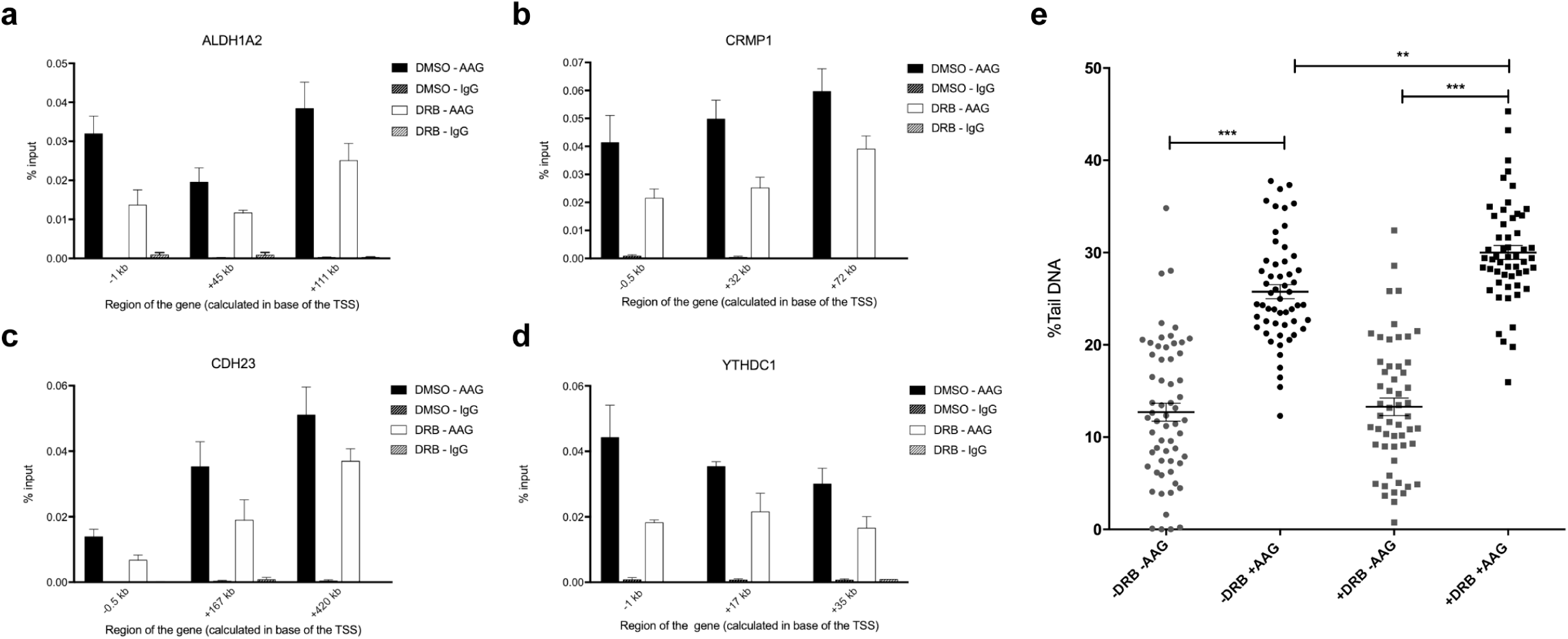
Transcription inhibition reduces AAG occupancy and causes accumulation of aberrant AAG substrates. **a-d** ChIP-qPCR experiments comparing AAG occupancy in DMSO and DRB treated HEK293T WT cells at *ALDH1A2* (a), *CRMP1* (b), *CDH23* (c) and *YTHDC1* (d) genes. Error bars, SEM from three independent experiments. **e** AAG Comet-FLARE assay in DMSO or DRB treated HEK293T cells. Error bars, SEM from at least three independent experiments. *p<0.05, **p<0.005, ***p<0.0005, two-way ANOVA.

### Loss of functional Elongator impairs AAG and APE1 chromatin recruitment and hinders AAG-initiated repair

Since AAG directly interacts with ELP1 and the two proteins co-occupy the same gene regions, we next addressed the importance of ELP1 for the recruitment of AAG-initiated BER to the chromatin. AAG ChIP in WT and ELP1^−/-^ cells revealed that loss of ELP1 causes global reduction in AAG binding to the chromatin at all genes tested (Fig. 4a-d). This effect was not a consequence of perturbed chromatin organization in ELP1^−/-^ cells, since histone occupancy was comparable in WT and ELP1^−/-^ cells, as indicated by histone H3 ChIP (Supplementary figure 7). Similar to the impact on AAG localization, loss of ELP1 resulted in a notable reduction of APE1 recruitment to all tested genes (Fig. 4e). Since ELP1 regulates AAG recruitment, we next tested if ELP1 status could influence the complex formation between AAG and hyperphosphorylated RNA pol II. IP of AAG in WT and ELP1^−/-^ HEK293T cells showed that loss of ELP1 diminishes the interaction between AAG and active RNA pol II (Fig. 4f). These findings suggest that ELP1 plays an important role in associating AAG with active transcription. To further evaluate the extent to which impaired AAG recruitment and reduced association with active transcription influence repair of aberrant bases in ELP1^−/-^ cells, AAG-initiated BER capacity was analyzed by flow cytometry host cell reactivation assay (FC-HCR)^33^. Importantly, cells lacking ELP1 showed significantly decreased capacity to repair AAG substrate hypoxanthine (Hx), when compared to WT cells (Fig. 4g). Hx was chosen over 3meA and 7meG, since these aberrantly methylated AAG substrates are unstable, and thus cannot be incorporated at the specific site in the DNA repair construct. Taken together, our findings suggest that a functional Elongator complex is a prerequisite for efficient recruitment of AAG-initiated BER to the chromatin and for its association with actively transcribing RNA pol II.

### Active transcription promotes AAG occupancy and stimulates repair of aberrant bases

Our findings suggest that AAG-initiated BER is associated with transcription through direct interaction with ELP1, and that Elongator promotes AAG-initiated BER in actively transcribing genes (Fig. 2 and 4). It remains however unclear to which extent active transcription directly influences AAG occupancy, and what is its importance in the repair of aberrant bases. To determine if transcription inhibition affects AAG distribution, ChIP experiments were performed in untreated cells and cells exposed to 5,6-dichloro-1-beta-D-ribofuranosylbenzimidazole (DRB). Treatment with transcription inhibitor DRB results in repression of transcription elongation, as consequence of impaired RNA pol II CTD phosphorylation^34,35^, thus causing reduced RNA pol II occupancy (Supplementary figure 8). Similar to the ELP1 loss (Fig. 4), transcription inhibition resulted in globally reduced AAG binding at all analyzed chromatin regions in DRB treated cells (Fig. 5a-d). This further suggests that transcription is an important modulator of AAG chromatin occupancy. To next determine if active transcription, besides promoting AAG occupancy, also has a role in AAG-initiated repair a single-cell AAG-Comet FLARE analysis was performed. Interestingly, inhibition of transcription elongation caused significant accumulation of AAG-specific aberrant bases, while it did not affect global DNA damage levels (Fig. 5e). In summary, these results strongly suggest that active transcription is required to promote AAG recruitment to the chromatin and to facilitate AAG-initiated BER of aberrant bases.

## Discussion

Removal and repair of aberrant bases are dramatically impaired in the context of chromatinized DNA^9-12,36^. It has been suggested that for efficient repair within chromatin to take place, BER needs to be associated with essential nuclear processes, such as transcription^6^. Besides the potential importance of transcription in promoting efficient BER, several studies indicated that BER enzymes, in particular DNA glycosylases, could influence transcription and play an important role in modulation of gene expression ^18-20,37^. However, direct evidence for the existence of transcription associated BER has not been provided so far. In this work we show that AAG-initiated BER associates with the transcription machinery, primarily by binding to the Elongator complex. Set of IP experiments indicates that the AAG DNA glycosylase forms a complex with active RNA pol II (Fig. 1C), predominantly through direct association with the ELP1 subunit of the Elongator (Fig. 2 and 4f). Identification of direct interaction between AAG and the ELP1 confirms recent predictions arising from a large proteomics study^21^. The relevance of this interaction is demonstrated by the RNA sequencing, presented in Fig. 2i, which revealed that AAG and ELP1 co-regulate a specific set of genes. On the chromatin level, the occupancy of the BER components AAG and APE1 accompanies the progressive enrichment of ELP1 and elongating RNA pol II at the 3’end of the co-regulated genes. The ELP1 and RNA pol II distribution patterns are strongly in line with the previous observations^29^. Importantly, regions of the co-regulated genes, enriched for AAG, APE1, ELP1 and RNA pol II S2P, present with the highest level of endogenous aberrantly methylated AAG substrates (Fig. 3m-p). Thus suggesting potential role of aberrantly methylated bases in regulation of gene expression. This is particularly interesting in light of recent work that indicated coevolution of epigenetic DNA methylation with repair of aberrantly methylated DNA bases, including AAG-initiated BER^38^. Increased level of aberrantly methylated AAG substrates at the 3’end of co-regulated genes (Supplementary figure 6) could directly be cause by RNA pol II accumulation, which indicative of a reduced elongation rate due to increased nucleosome density^39^. This could directly lead to reduced repair and consequent accumulation of aberrant bases. Indeed, the findings provided by genome-wide mapping of aberrantly methylated bases in yeast, suggested that repair of AAG substrates is slower in the regions with strongly positioned nucleosomes, in comparison to the nucleosome depleted regions^40^.

As in the case of RNA pol II complex formation, the recruitment of AAG-initiated BER to the chromatin is strongly dependent on the functional Elongator, and results in dramatic decrease of AAG and APE1 occupancy in ELP1^−/-^ cells (Fig. 4). Besides its impact on the chromatin recruitment analysis of AAG-repair capacity in ELP1^−/-^ cells revealed that Elongator has an important role in facilitating efficient AAG-initiated repair (Fig. 4g). This effect is directly dependent on active transcription as suggested by experiments involving transcription elongation inhibitor DRB. Notably, DRB treatment resulted in reduced AAG occupancy and impaired AAG-initiated repair (Fig. 5), thus suggesting that active transcription has an important role in the maintenance of genome stability by facilitating BER. Very recently Menoni et al. similarly indicated that the inhibition of transcription elongation by DRB results in reduced recruitment of the BER scaffold protein XRCC1 to the sites of initiated repair^41^. The fact that inhibition of transcription elongation impairs AAG-initiated BER and results in global accumulation of aberrantly methylated AAG substrates, strongly supports the idea that one key functional consequence of AAG association to the active transcription is to ensure efficient repair. Accumulation of AAG substrates genome wide could directly have harmful effects on transcription, since 3meA was shown to affect the fidelity of incorporation by RNA pol II^42^. While the 3meA analogue 3-deaza-3meA can successfully be bypassed by RNA pol II, its presence causes 10-fold greater misincorporation of CMP, thus resulting in synthesis of mRNA’s bearing mutations. In contrast to 3meA, AP site intermediates, generated by APE1 during BER, impair RNA pol II progression^43^. Finding that AAG, similarly to other DNA glycosylases^37^, primarily inhibits gene expression (Fig. 1g) suggests that by coordinating BER initiation with transcription, AAG could potentially inhibit RNA pol II progression and therefore repress generation of faulty transcripts. Imbalanced AAG-initiated BER and consequent accumulation of aberrantly methylated bases can thus, besides causing genome instability and increased mutation rate, also impact gene expression.

Loss of AAG results in increased expression of genes that primarily segregate in neurodevelopmental processes (Fig. 1 and Supplementary figure 1). It has been demonstrated recently that the DNA glycosylases Ogg1 and Mutyh, involved in repair of oxidized bases, play an important role in neurodevelopment through impact on transcription. Lack of *Ogg1* and *Mutyh* was reported to up-regulate ERα target-genes, which in turn modulated cognition and anxiety-like behavior in mice^20^. It will be thus interesting to determine the importance of transcription associated AAG-initiated BER in regulation of neurodevelopment, and test its role in brain functioning.

Taken together, our results suggest that the transcriptional status, in addition to the type of aberrant bases and DNA glycosylases involved, is an essential determinant of BER efficiency and its impact on genome and chromatin organization. Association of AAG with transcription elongation could thus, in addition to enabling efficient removal of aberrantly methylated bases, serve as an important novel layer of gene expression control.

## Methods

### Cells and culturing

Human embryonic kidney cells expressing a mutant version of the SV40 large T antigen (HEK293T) were maintained under 5% CO_2_ and 37 °C in DMEM high glucose supplemented with 10% fetal bovine serum, and 1% penicillin/streptomycin.

### Fractionation

To obtain total fraction (TF) 2.5 × 10^6^ HEK293T cells were resuspended in 200 µL of MNase buffer (50 mM Tris-HCl pH 7.5, 30 mM KCl, 7.5 mM NaCl, 4 mM MgCl_2_, 1 mM CaCl_2_, 0.125% (v/v) NP-40, 0.25% Na-deoxycholate, 0.3 M sucrose, 1x Halt™ Protease Inhibitor Cocktail) with 1U MNase, and the samples incubated at 37°C for 30 min. For the preparation of the soluble fraction (SF) and chromatin fraction (CF) 2.5 × 10^6^ cells were resuspended in 200 µL of chromatin extraction buffer (10 mM HEPES pH 7.6, 3 mM MgCl_2_, 1 mM DTT, 1x Halt™ Protease Inhibitor Cocktail) and samples rotated 30 min at RT. SF was separated from nuclei by low-speed centrifugation (1,300*g* for 10min) and collected for further analysis. Pelleted nuclei were lysed in 200 µL of MNase buffer with 1U MNase, for 30 min at 37°C. TF, SF and CF were boiled in Laemmli buffer, centrifuged at 16,000*g* for 5 min at RT, and equal volumes of each fraction subjected to immmunoblott analysis.

### Generation of HEK293T AAG^−/-^, ELP1^−/-^, AAG^−/-^ ELP1^−/-^ and HA-ELP1 cell lines

CRISPR short guide RNAs (sgRNAs) were designed using the Optimized CRISPR Design tool (http://tools.genome-engineering.org). The oligo pairs encoding the sgRNAs (Supplementary table 4) were annealed, ligated into pSpCas9(BB)-2A-GFP (PX458)4 (Addgene plasmid # 48138; a gift from Feng Zhang) and knockout (KO) cell lines generated according to^44^. Mutations and loss of target protein expression was confirmed in the clonal KO cell lines by sequencing and immunoblot analysis, respectively. The presence of off-target mutations in KO cells was excluded by sequencing of top-candidate off-target sites predicted by Optimized CRISPR Design tool. For generation of HEK293T HA-ELP1 cells plasmid containing the sgRNA for ELP1 C-terminus was transfected together with the HA-ELP1 repair oligo containing BspEI specific cut-site (Supplementary table 4) into HEK293T cells using nucleofector (Lonza), according to the manufacturer’s protocol. The cells were tested for HA insertion by: PCR followed by specific BspEI digestion, sequencing and immunoblot analysis.

### RNA isolation, library preparation and RNA sequencing

Total RNA was purified with RNeasy kit (QIAGEN), and DNAseI digested (QIAGEN) according to the manufacturer’s protocol. RNA library preparation and sequencing were carried out by the Functional Genomic Centre, Zurich (FGCZ). The libraries were screened on the Bioanalyzer (Agilent), pooled in equimolar concentrations, and sequenced on one lane of a flow cell on an Illumina HiSeq2000 using single-read TruSeq v3 chemistry with 125-cycles.

### Bioinformatic Processing

FASTQ files were processed to remove adapter sequences and low quality reads prior to alignment to the GRCh37 Ensembl release 80 of the human genome using STAR v2.4.2a aligner^45^. Reads within exon features were counted using featureCounts in Bioconductor (3.2) package Rsubread (1.20.6). Differential expression analysis was performed in R using the Bioconductor package DESeq2^46^. Pairwise comparisons were made between the HEK293T WT and the knockout cell-lines, AAG^−/-^ or ELP1^−/-^ samples. For the WT vs AAG^−/-^ comparisons, statistically significant differentially expressed genes were defined as having a ≥1.5-fold change and FDR≤0.1. For the WT vs ELP1^−/-^ comparisons, statistically significant differentially expressed genes were defined as having a ≥2-fold change and FDR≤0.1. DAVID^31^ gene-ontology (GO) enrichment analysis was performed to identify enriched Biological Processes (BP).

### Protein mass spectrometry

FLAG-tagged full-length human AAG was expressed and purified from 1h mock or 2 mM MMS treated HEK293T cells as described previously^8^. Identification of protein complexes in FLAG-AAG and control FLAG IP samples was performed by the Biopolymers and Proteomics Core Facility of the David H. Koch Institute for Integrative Cancer Research (http://ki.mit.edu/sbc/biopolymers). Briefly, protein samples were reduced, alkylated, and digested with trypsin, followed by purification and desalting on an analytical C18 column tip. The processed peptides were then analyzed on an Agilent Model 1100 Nanoflow high-pressure liquid chromatography (HPLC) system coupled with electrospray ionization on a Thermo Electron Model LTQ Ion Trap mass spectrometer (MS). All MS/MS samples were analyzed using Sequest (Thermo Fisher Scientific, San Jose, CA, USA; version SRF v. 3). Sequest was set up to search the merged_human_90T.fasta.hdr database assuming the digestion enzyme trypsin. Sequest was searched with a fragment ion mass tolerance of 0.50 Da and a parent ion tolerance of 2.0 Da. Dehydro of serine and threonine, oxidation of methionine, iodoacetamide derivative of cysteine and phosphorylation of serine, threonine and tyrosine were specified in Sequest as variable modifications. Scaffold (version Scaffold_4.3.0, Proteome Software Inc., Portland, OR) was used to validate MS/MS based peptide and protein identifications. Peptide identifications were accepted, if they could be established at greater than 95.0% probability (as specified by the Peptide Prophet algorithm^47^). Peptide identifications were also required to exceed specific database search engine thresholds. Sequest identifications required at least deltaCn scores of greater than 0.10 and XCorr scores of greater than 2.0, 2.5, 3.5 and 3.5 for singly, doubly, triply and quadruply charged peptides. Protein identifications were accepted if they could be established at greater than 99.0% probability and contained at least 2 identified peptides. Protein probabilities were assigned by the Protein Prophet algorithm^48^. Proteins that contained similar peptides and could not be differentiated based on MS/MS analysis alone were grouped to satisfy the principles of parsimony. All proteins identified in Flag-AAG and control Flag samples, of cells untreated or treated with MMS are indicated in Table S1 and available in PRIDE under accession PXD013508.

### Whole cell extracts

HEK293T cells were trypsinized and washed twice with ice cold PBS and the pellet flash frozen in liquid N_2_. Whole cell extracts (WCEs) were prepared as described previously^8^. In case of nuclease treatment either 1.2 µl CaCl_2_ (2.5 M) with 25U of DNAse, or 2.1 µg RNAseI, and 50U MNase and samples incubated for 1h, at 4 °C. Samples were centrifuged and the WCE supernatant collected.

### Co-immunoprecipitation (CoIP) assays

WCEs (0.5 mg) were incubated with 2 µg of antibody (anti-AAG antibody^8^; or anti-ELP1, Abcam ab56362) or IgG (rabbit IgG, Diagenode, C15410206; or mouse IgG, Diagenode, C15400001) at 4°C overnight. In case of purified proteins (FLAG-ELP1, fl-AAG or Δ80 AAG) 1µg of protein was mixed with 1µg of antibody and incubated overnight at 4 °C in 200 µl final volume of IP buffer (20 mM HEPES pH 7.9, 2 mM MgCl_2_, 0.2 mM EGTA, 10% (v/v) glycerol, 0.1 mM PMSF, 2 mM DTT, 140 mM NaCl, 0.01% (v/v) Nonidet-P40). Protein-A Dynabeads, or Protein-A sepharose beads (in case of purified proteins), were equilibrated in the IP buffer at 4 °C overnight. After three consecutive washes the beads were added to samples and incubated for 4 h at 4 °C. The supernatant was removed and beads washed three times with cold wash buffer. Beads were boiled in Laemmli buffer and subjected to immunoblot analysis.

### Immunoblot analysis

Samples were separated on 4–12% Bis–Tris polyacrylamide gel (Invitrogen) followed by transfer to Amersham™ Hybond^®^ P 0.2 PVDF (GE Healthcare) for immunoblotting. Primary antibodies: anti-anti-ELP1 (Abcam, ab56362), anti-ELP3 (Abcam, ab96781), anti-AAG (described in ^8^), anti-RNA pol II S2P (Abcam, ab5098), anti-HA (Abcam, ab9110), anti-α-Tubulin (Cell Signaling, 2144); were detected using infrared (IR) Dye-conjugated secondary antibodies (Li-COR Bioscienecs, 827-11081 and 925-32210). The signal was visualized by using direct IR fluorescence via the Odyssey Scanner, LI-COR Biosciences.

### Column purification of cellular complexes

Cellular complexes were isolated from HeLa cell pellets previous published protocol^22^. Briefly 1ml, HeLa cell pellet was washed twice with 1 packed cell volume (PCV) of ice-cold PBS, and resuspended in 4PCV of hypotonic buffer and incubated at 0°C for 20 min. After addition of protease inhibitors, cells were homogenized using a Douncer and 4PCV of ice-cold buffer 2 were slowly added while stirring. Next, 1 PCV of neutralized saturated ammonium sulfate solution was added during stirring for 30 min at 0°C, followed by ultracentrifugation 3 h at 2°C, 45’000 rpm. The supernatant was collected and dialyzed into buffer B with 50 mM KCl; 3 times for 1h at 4°C. Dialized samples were load onto heparin-sepharose column overnight. After washing with 10 column volumes of buffer B supplemented with 50 mM KCl, bound proteins were eluted as 1mL fractions using buffer B with 150 mM, 300 mM, 450 mM, 700 mM and 1000 mM KCl, sequentially. The different elutions were subjected to immunoblot analysis.

### Protein purification

Human fl AAG and Δ80AAG lacking first 80 N-terminal amino acids were expressed and purified as described previously^49^. ELP1 cDNA was excised from pCMV6-XL5-ELP1 (OriGene) by NotI (NEB) and inserted into pcDNA3.1(+)-3xFLAG vector generated as described in^8^. HEK293T cells were transfected by calcium phosphate, harvested after 48 h and WCE prepared as described above. 0.5 mg WCE were incubated with 75 µL anti-FLAG M2 affinity beads (Sigma Aldrich) for 2 h at 4 °C, the beads were washed three times with washing buffer (20 mM Hepes pH 7.9, 2 mM MgCl2, 0.2 mM EGTA, 10 % glycerol, 0.1 mM PMSF, 2 mM DTT, 1x Halt™ Protease Inhibitor Cocktail). FLAG-Elp1 was eluted twice by addition of 80 µL Elution buffer (washing buffer with 0.5 % NP-40 and 0.15 µg/µL 3x FLAG peptide (Sigma Aldrich)) for 30 min. The supernatant was applied on an Amicon Ultra-0.5 Centrifugal Filter Unit (Merck) and buffer exchanged to storage buffer (washing buffer with 0.1 % NP-40).

### Gene expression analysis

RNA was purified with RNeasy kit (QIAGEN) according to the manufacturer’s protocol and reverse transcribed using MultiScribe Reverse Transcriptase (QIAGEN) according to the manufacturers protocol. qPCR was performed with Power SYBR Green PCR Master Mix (Applied Biosystems) on a StepOnePlus Real-Time PCR System. Relative transcription levels were determined by normalizing to GAPDH mRNA levels. All primer sequences are listed in Supplementary table 2.

### Chromatin immunoprecipitation

HEK293T cells were crosslinked with 1% formaldehyde and quenched with 0.110 mM glycine final. Cells were washed with ice-cold PBS, and harvested. Cell pellets were resuspended in cell lysis buffer (100mM Tris-HCl pH 8, 10 mM DTT) and incubated 15min on ice, and 15min at 30°C shacking. Next, nuclei were pelleted and washed with buffer A (10 mM EDTA pH 8, 10 mM EGTA, 10 mM HEPES pH 8, 0.25% Triton X-100), followed by buffer B (10 mM EDTA pH 8, 0.5 mM EGTA, 10 mM HEPES pH 8, 200 mM NaCl). Nuclei were lysed in lysis buffer (50 mM Tris-HCl pH 8, 10 mM EDTA and 1% SDS) and chromatin sheered to 200-250bp DNA fragments by sonication with Bioruptor (Diagenode) for 30 cycles of 30 seconds. 40 µg of chromatin was next precleared 2h at 4°C, and then incubated with 2 µg antibody in ChIP buffer (16.7 mM Tris-HCl pH 8, 167 mM NaCl, 1.2 mM EDTA, 0.01% SDS and 1.1% Triton X-100) overnight at 4°C. ChIP antibodies included: anti-AAG (LSBio, LS-C133325-100); ant-HA (Abcam, ab9110); anti-APE1 (Abcam, ab194); anti-RNA pol II S2P (Abcam, ab5098); anti-H3 (Abcam, ab1791); anti-RNA pol II (MBL, MABI0601). The DNA-protein-antibody complexes were isolated using A/G dynabeads (Thermofischer) and washed using sequentially: low salt wash buffer (16.7 mM Tris-HCl pH 8, 167 mM NaCl, 0.1% SDS, 1% Triton X), high salt wash buffer (16.7 mM Tris-HCl pH 8, 500 mM NaCl 0.1% SDS, 1% Triton X) and LiCl wash buffer (250 mM LiCl, 0.5% NP40, 0.5% Na-deoxycholate, 1mM EDTA, 10mM Tris-HCl pH 8). Proteinase K treatment was performed for 1 hr at 50°C with 10 mM EDTA, 40 mM Tris-HCl pH 6.5 and 20 µg proteinase K. The DNA was purified with phenol-chloroform, ethanol precipitated and analyzed by qPCR. Levels of ChIPed DNA is expressed as % of input, or relative abundance = (% input of specific gene region) / (% input of promoter region). Primer sequences are listed in the Supplementary table 3.

### Real-time qPCR-based assay for quantification of aberrant AAG substrates

The levels of aberrant bases substrates of AAG were determined in sheared HEK293T WT genomic DNA with an average fragment size of 200bp. DNA was incubated with either a combination of 10U of AAG (NEB, M0313) and 10U of APE1 (NEB, M0282), or only with 10U of APE1 for 1h at 37°C in 1X ThermoPol Reaction Buffer (NEB, B9004). Next, level of DNA damage was assessed by qPCR using primers targeting regions corresponding to promoter, middle and end of the gene of interest (Supplementary table 3). Level of AAG substrates in the targeted regions was inferred by calculating: ΔCt = Ct(+AAG+APE1) - Ct(-AAG+APE1). The relative amount of AAG substrates for each gene region was calculated with respect to the promoter.

### Flow cytometry host cell reactivation assay (FM-HCR)

FM-HCR assay was performed as previously described^33^, using plasmids for expression of the fluorescent proteins EGFP and mPlum subcloned into the pmaxCloning. EGFP plasmid served as control. The repair mPlum plasmid was engineered by placing the AAG-specific base lesion hypoxanthine (Hx) lesion into the open reading frame of *mPlum*. In case of inefficient repair, Hx is maintained in the plasmid and due to transcriptional mutagenesis at the site of lesion results in the expression of a non-fluorescent reporter protein. Fluorescent mPlum is generated only upon efficient BER and removal of the base lesion. The mPlum fluorescence is thus proportional to the level of AAG-initiated BER. During flow cytometry analysis, the cell debris, doublets, and aggregates were excluded based on their side-scatter and forward-scatter properties. To exclude dead cells Zombie NIR (BioLegend) dye was added to cells 15–30 min before the analysis. The following fluorophores and corresponding detectors (in parentheses) were used: EGFP (Alexa Fluor 488), mPlum (PE-Cy5-5), and Zombie NIR (Alexa Fluor 700). Calculation of Percent Fluorescent Reporter Expression %R.E. was computed as previously described^33^.

### Comet FLARE

The level of AAG-specific substrates in actively transcribing and inhibited cells was assessed by single-cell AAG-Comet FLARE (Fragment Length Analysis using Repair Enzymes) assay. Briefly, transcriptional stalling was induced by treatment of HEK293T WT cells with 150 μM DRB for 4 h at 37 °C, DMSO served as control. Next cells were collected and mixed with 1% of low melting point agarose, spread onto glass slides (Trevigen) and lysed overnight in lysis buffer (2.5 M NaCl, 100 mM EDTA, 10 mM Tris, 1% Triton, 10% DMSO, 1% sodium lauryl sarcosinate). Next, detergent was removed by washing and the slides were incubated for 1h at 37°C in the presence of 2U of AAG (NEB, M0313) in reaction buffer (2.5 mM HEPES-KOH pH 7.4, 100 mM KCl, 25 mM EDTA, 2 mM BSA). The DNA was subjected to alkaline electrophoresis (1 mM EDTA, 200mM NaOH, pH > 13) and labelled with SYBR^®^ Green (Sigma Aldrich). In total, four replicates were performed and one hundred cells (fifty cells per technical duplicate) analyzed per experiment. The percentage of the DNA in the tail was evaluated for each data point with Comet IV, version 4.2 (Perceptive Instruments, Suffolk, U.K.).

### Quantification and Statistical Analysis

Immunoblots were quantified by GelEval 1.35 scientific imaging software (FrogDance Software, UK). Analysis of data was performed using GraphPad Prism (GraphPad Software, Inc., La Jolla, CA). Statistical significance was determined primarily by two-tailed unpaired Student’s t-test. All data represent mean values ± SEM; *p<0.05; **p<0.01; ***p<0.001 versus control; NS, not significant. Comet FLARE was analyzed using two-way ANOVA. All data represent mean values ± SEM. *p<0.05; **p<0.01; ***p<0.001 versus control; NS, not significant.

### Data Resources

The RNA sequencing data reported in this paper are available in GEO under accession GSE129009 for HEK293T cells and GSE129010 for HAP1 cells. The proteomics data associated with Fig. 2 and Supplementary table 1 are available in PRIDE under accession PXD013508.

## Acknowledgements

This work has been supported by the Norwegian Research Council grant (ID: 263152/F20) to B.v.L and N.P.M.; Onsager Fellowship to B.v.L; NTNU to D.L.B. and S.B.; University of Zurich to M.R. and A.F. We thank to Raffaella Santoro, Eva Vollenweider and Sergio Leone for help with initial ChIP experiments and RNA-sequencing analysis; to Hilde Nilsen, Lisa Lirussi and Alexander Chaim for suggestions and shearing protocols and reagents; to Cathrine Broberg Vågbø and PROMEC for analysis of base modifications; to Hans E. Krokan, Katja Scheffler and Cathrine Broberg Vågbø, for discussions and comments to the manuscript; and to FGCZ for RNA-sequencing.

## Author Contributions

B.v.L. and L.D.S. had the original idea. B.v.L. supervised the project and together with N.P.M. and M.B. interpreted the experiments. B.v.L., N.P.M. and D.L.B. wrote the manuscript and generated figures. N.P.M. performed most of the experiments D.L.B. performed the qPCR analysis of AAG substrates and Comet-FLARE. A.B. contributed to ChIP and IP experiments. M.R. purified proteins and contributed to IP studies. S.B. contributed to qPCR analysis. K.Ø.B and M.O. contributed with RNA sequencing of HAP1 cells. P.A.A. contributed to generation of HA-ELP1 cells. A.F. performed isolation of complexes. S.F.M., L.C.O. and P.S. contributed to bioinformatics analysis.

