## Supplementary Information for "“Alkyladenine DNA glycosylase associates with transcription elongation to coordinate DNA repair with gene expression”"

**SUPPLEMENTARY TABLES**

**Table S1.** Proteomics Data for FLAG and FLAG-AAG Co-immunoprecipitation.

**Table S2.** List of DNA primers used for RT-qPCR analysis of mRNA level.

| **cDNA primers** | | |
|---|---|---|
| **Name** | **Forward / Reverse** | **Sequence 5' to 3'** |
| GAPDH | Fw | GAGTCAACGGATTTGGTCGT |
| | Rv | TTGATTTTGGAGGGATCTCG |
| ALDH1A2 | Fw | GCTCTCATGGTATCCTCCGC |
| | Rv | TCCCGTAAGCCAAACTCACC |
| CRMP1 | Fw | CTCTGCTGTCAGTTCGCCAG |
| | Rv | GATGCGTCCACCTCTGATGA |
| CDH23 | Fw | CGGAGTGGATCCCAGAGAGA |
| | Rv | GCAAGTGACCAGGGAGTACC |
| YTHDC1 | Fw | GAGGGCCAAATCTCCTACGC |
| | Rv | GTCTCATGGTCAGAGCCATATTC |
| SYT9 | Fw | GGGATCCAGGAGAACTGTGC |
| | Rv | TCCGGGTTTGACAAGTTGAGT |
| CDH4 | Fw | GATGTACGTCACAAGGCCCA |
| | Rv | AGGGCGGTTGTCATTCATGT |
| NPTX2 | Fw | TCTGGTGACTTAAAGGCGCT |
| | Rv | CTGCACAATGAGACCTCGG |
| NOVA2 | Fw | ACTGTTCCATCACGGCTTTC |
| | Rv | ACGATGAACCCCGACAGA |
| CDH22 | Fw | CACAACATCACAGTGCTGGC |
| | Rv | CATACAGCTGCCTCGTAGGG |

**Table S3.** List of DNA primers used for ChIP analysis.

| **ChIP primers** | | |
|---|---|---|
| **Name** | **Forward / Reverse** | **Sequence 5' to 3'** |
| ALDH1A2 -0.5 kb | Fw | GAAGTCGAGCGAGGGTCG |
| | Rv | CTCTACTCCGAAGCAGCACC |
| ALDH1A2 +45 kb | Fw | CTGACAGAATTTATCTGAAGGACTC |
| | Rv | ATGAGGTAAAACTGACCATAGAAGC |
| ALDH1A2 +111 kb | Fw | GGATGGGAAGAAAGGGAGGC |
| | Rv | GCCAAGTTCCATTGTGCCAG |
| CRMP1 -0.5 kb | Fw | GTGGTAAGGGCCAGGAAAGG |
| | Rv | TGTGTCTGTCGGAGTGTGC |
| CRMP1 +32 kb | Fw | AGAAGTCATCAGCCGCAGTC |
| | Rv | CGGATGGTTATTCCCGGAGG |
| CRMP1 +72 kb | Fw | CAGGTCAGTGGGCAGTTCAT |
| | Rv | GTTGACGGGGCAGTCAGATT |
| CDH23 -0.5 kb | Fw | GTTGAGGCCCAGAAAGTCTCA |
| | Rv | TTCCGAGCTGTCACTGTTCC |
| CDH23 +167 kb | Fw | CTCCTTGCCAGGCCCATTTA |
| | Rv | TATGTTTCATGCCCTGCGGT |
| CDH23 +420 kb | Fw | CCCCACGTGGACAAGAAAGT |
| | Rv | AACCGGAGGGAACATCAGTC |
| YTHDC1 -1 kb | Fw | AGCAGACCATCGAGGAATCG |
| | Rv | TGTTTCCGTTGGACAATGAATCT |
| YTHDC1 +17 kb | Fw | AGCCATTATCACACAAAGGGGT |
| | Rv | AGCCTTCCAGGTTGAATGGG |
| YTHDC1 +35 kb | Fw | ATCGGCGGAGAGAACAATCC |
| | Rv | ACCATGTCAGCATATTACCTTCTGT |

**Table S4.** List of DNA oligonucleotides used for CRISPR-Cas9 editing.

| DNA oligonucleotides | Sequence 5' to 3' |
|---|---|
| $AAG^{-/-}$ sgRNA | GCTGGCGGTGCCTTTCCGGA |
| $AAG^{-/-}$ sgRNA top | CACCGGCTGGCGGTGCCTTTCCGGA |
| $AAG^{-/-}$ sgRNA bottom | AAACTCCGGAAAGGCACCGCCAGCC |
| $ELP1^{-/-}$ sgRNA | AGCTCAGATTCGGAAGTGGT |
| $ELP1^{-/-}$ sgRNA top | CACCGAGCTCAGATTCGGAAGTGGT |
| $ELP1^{-/-}$ sgRNA bottom | AAACACCACTTCCGAATCTGAGCAC |
| HA-ELP1 sgRNA | GACTGAGTGACTGCAGTTAGG |
| HA-ELP1 sgRNA top | CACCGACTGAGTGACTGCAGTTAGG |
| HA-ELP1 sgRNA bottom | AAACCCTAACTGCAGTCACTCAGT |
| PCR #1 HA insertion Fw: | TTGCTCTGACCTGGAGGTTG |
| PCR #1 HA insertion Rv: | GTAGTCCGGAACGTCGTAG |
| PCR #2 BspEI digestion Fw: | TTGCTCTGACCTGGAGGTTG |
| PCR #2 BspEI digestion Rv: | GCTCTCAAACAGCCCAAGTG |
| HA-ELP1 repair oligo: | TTGCTTTTAGATGCTGAGCTTTTTATACCACCAAAGATC AACAGAAGAACCCAG TGGAAGCTGAGCCTGCTAGACTACCCCTACGACGTTCC GGACTACGCCTGAGAGAAGACCATTTCCACTCATTCCT GTTGTCCTACCACCCCTTGCTCTTTGAGGGCTGGCTAT TGAGAACTGGAA |

**SUPPLEMENTARY FIGURES**

**Supplementary figure 1.** Loss of AAG alters expression of neurodevelopmental genes in HAP1 cells. **a** Immunoblot of whole cell extracts from HAP1 WT and $AAG^{-/-}$ cell lines generated via CRISPR-Cas9 technology. **b** Top six biological processes (BP) gene ontology (GO) terms as determined by the Database for Annotation, Visualization and Integrated Discovery (DAVID) for genes dysregulated in $AAG^{-/-}$ in HAP1 cells.

**Supplementary figure 2.** AAG and ELP1 subunit of transcriptional Elongator complex directly interact. **a** Separation of cellular complexes from HeLa cells by heparin-spharose affinity chromatography. The elutions at the indicated potassium chloride (KCl) concentrations were immunoblotted and probed for AAG and ELP1 and ELP3. **b** AAG-mediated immunoprecipitation of whole cell extracts from HEK293T treated and untreated DNAseI. **c** IP of AAG from HEK293T WCEs untreated or treated with DNAseI, Mnase and RNAseI. **d** Schematic representation of full-length (fl) AAG and AAG lacking 80 N-terminal amino acids (Δ80); numbers indicate amino acids. **e-g** SDS-PAGE analysis of purified recombinant Δ80 AAG (e); fl AAG (f); and FLAG-tagged ELP1 (g).

**Supplementary figure 3.** AAG and ELP1 regulate expression of neurodevelopmental genes**. a-e** Expression of additional neurodevelopmental genes *SYT9* (a), *CDH4* (b), *NPTX2* (c), *NOVA2* (d), *CDH22* (e) in WT, $AAG^{-/-}$, $ELP1^{-/-}$ and $AAG^{-/-}$ $ELP1^{-/-}$. Error bars, SEM from at least three independent experiments. *p<0.05, **p<0.005, ***p<0.0005, two-tailed Student's t test; NS, not significant.

**Supplementary figure 4.** AAG regulatory effect on expression of neurodevelopmental genes is specific and reproducible in different knockout clones. **a** Immunoblot of whole cell extracts of HEK293T WT, $AAG^{-/-}$ clone A and $AAG^{-/-}$ clone B generated via CRISPR-Cas9 technology. **b-j** mRNA expression levels of *ALDH1A2* (b), *CRMP1* (c), *CDH23* (d), *SYT9* (e), *CDH4* (f), *NPTX2* (g), *NOVA2* (h), *CDH22* (i) and *YTHDC1* (j) genes in WT, $AAG^{-/-}$ clone A and $AAG^{-/-}$ clone B. Error bars, SEM from at least three independent experiments. *p<0.05, **p<0.005, ***p<0.0005, two-tailed Student's t test.

**Supplementary figure 5.** Generation of HA-ELP1 HEK293T cell lines. **a** Schematic representation of the used approach. **b** Identification of the clone homozygous for the HA insertion, by PCR and subsequent enzymatic digestion with BspEI. **c** Immunoblot of whole cell extracts of HEK293T WT, $AAG^{-/-}$, $ELP1^{-/-}$ and WT HA-ELP1 (heterozygous and homozygous) cell lines generated via CRISPR-Cas9 technology. The HA-ELP1 cell lines tested by immunoblotting were additionally sequenced confirming the HA insertion in frame with ELP1.

**Supplementary figure 6.** RNA polymerase II distribution along co-regulated genes. **a-c** ChIP assays showing relative RNA Pol II S2P (RNA pol II phosphorylated at Serine 2 of the C-terminal domain) occupancy in genes co-regulated by AAG and ELP1: *CRMP1* (a), *CDH23* (b) and unaffected gene *YTHDC1* (c) in WT HEK293T cells. Values are shown as relative occupancy: % input of specific gene region relative to % input of promoter region Error bars represent the SEM calculated from three biological replicates. *p<0.05, **p<0.005, ***p<0.0005, two-tailed Student's t test.

**Supplementary figure 7.** Loss of ELP1 does not affect H3 occupancy**.** ChIP-qPCR experiments targeting histone 3 (H3) in promoters of co-regulated genes in WT and ELP1$^{-/-}$ HEK293T cells.

**Supplementary figure 8.** DRB treatment reduces RNA polymerase II occupancy. **a-c** ChIP-qPCR experiments comparing RNA polymerase II occupancy in DMSO and DRB treated WTCas9 HEK293T cells at *CRMP1*(a), *CDH23* (b) and *YTHDC1* (c) genes.

### Supplementary figure 1

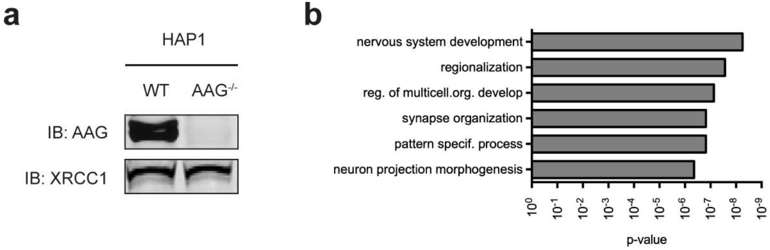

### Supplementary figure 2

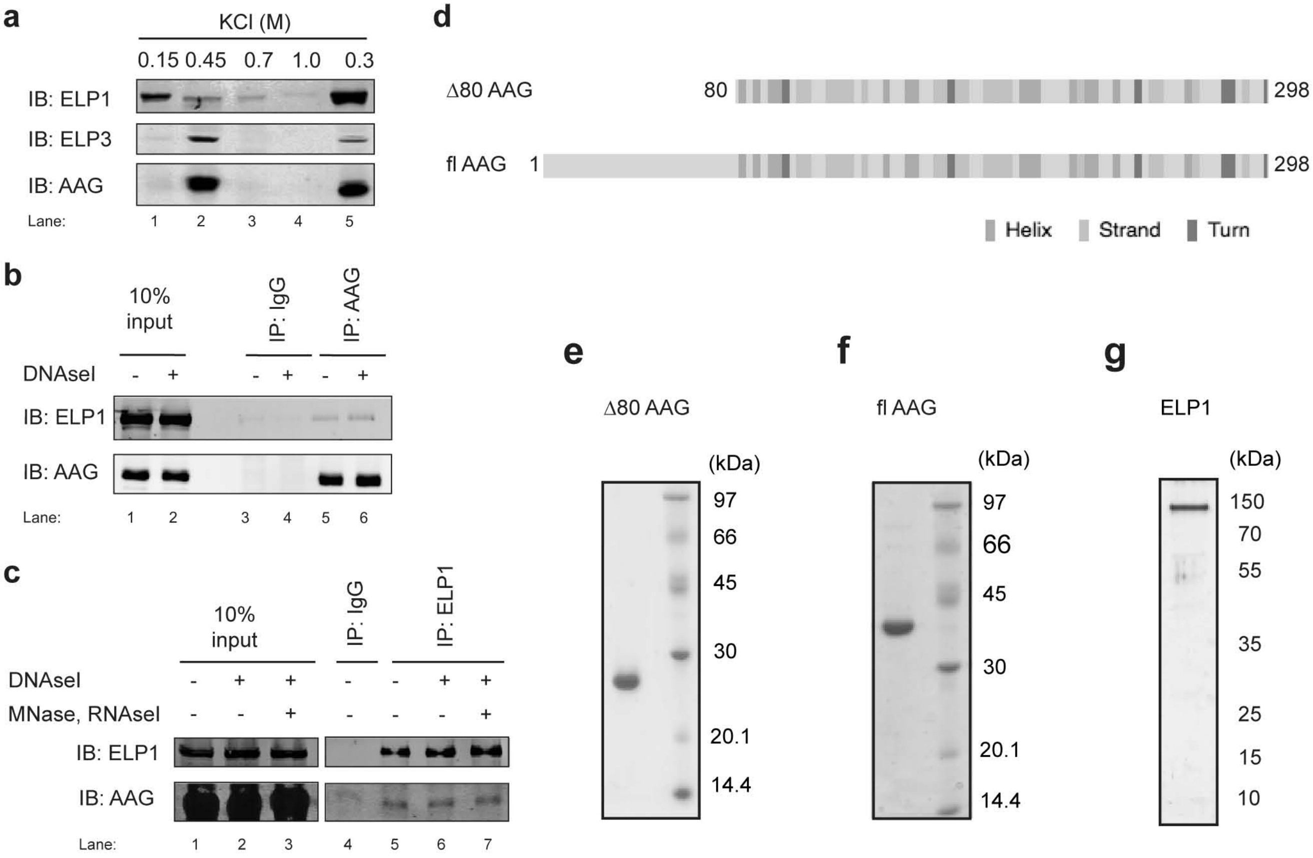

### Supplementary figure 3

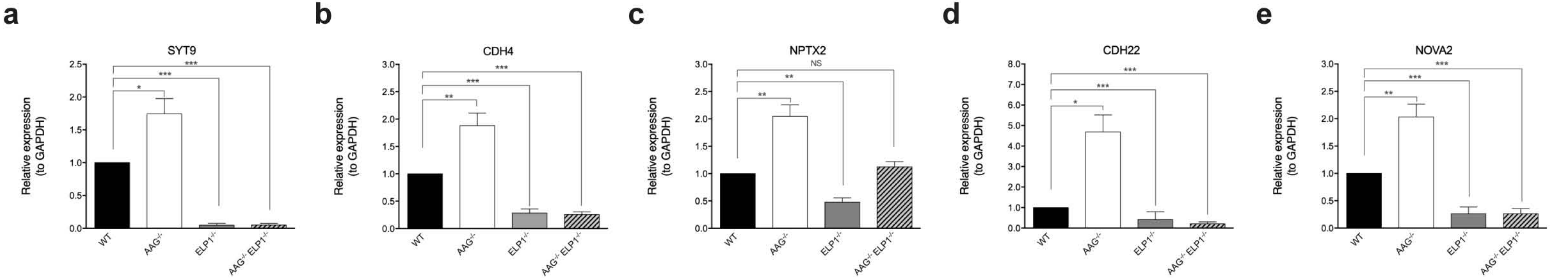

### Supplementary figure 4

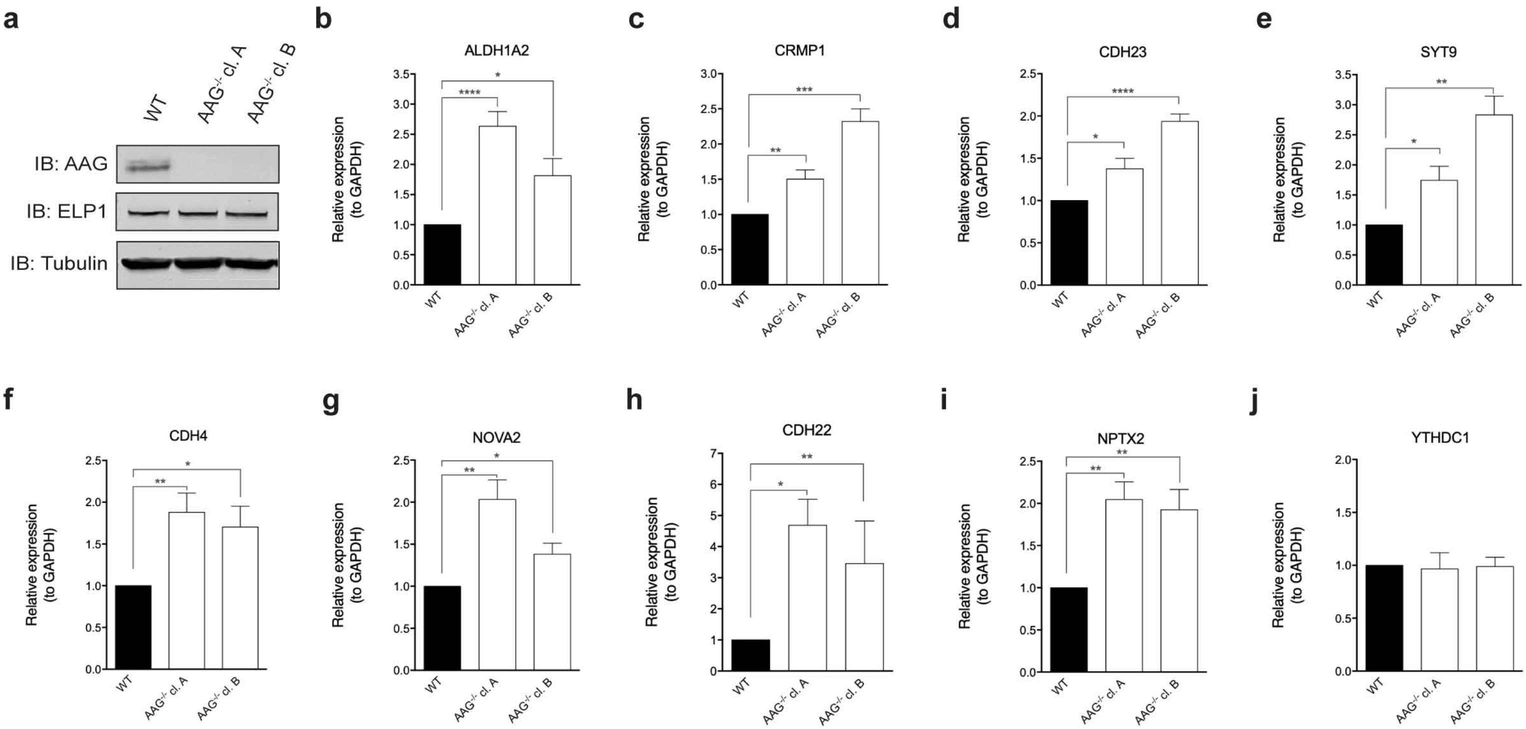

### Supplementary figure 5

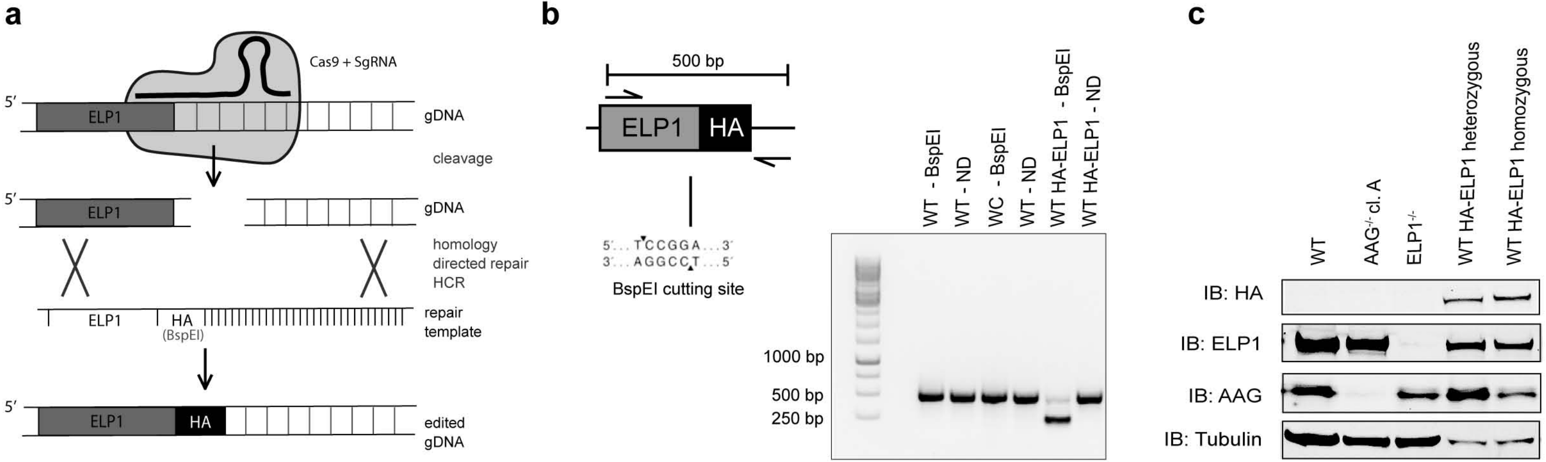

### Supplementary figure 6

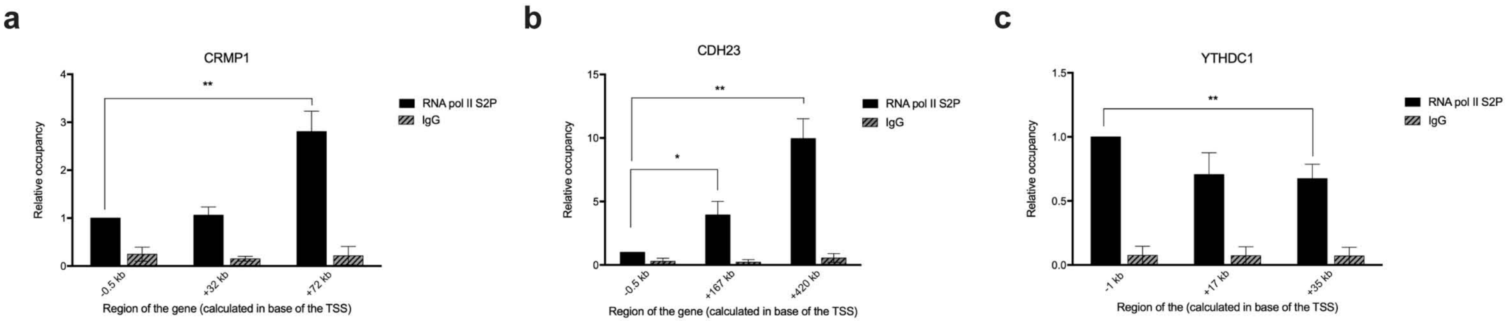

### Supplementary figure 7

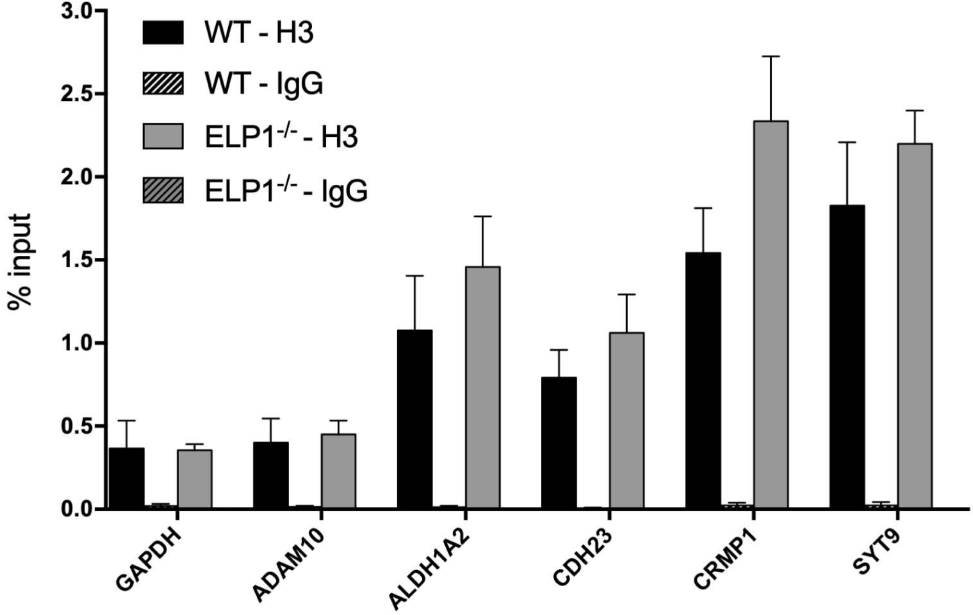

### Supplementary figure 8

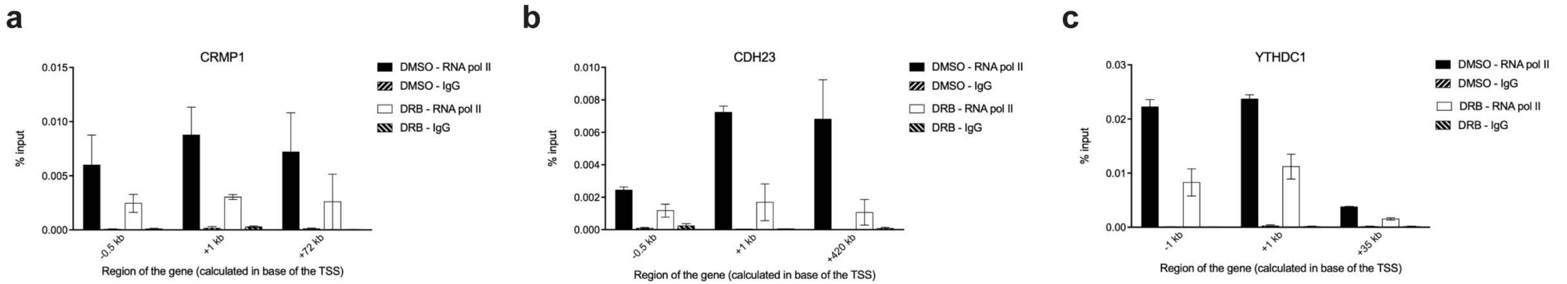
